# Downstream mRNA Secondary Structure, Not Codon Elongation Supply, Coordinates Co-Translational Protein Folding Across the Human Ribosomal Exit Tunnel

**DOI:** 10.64898/2026.09.27.753847

**Authors:** Muhammad Saad Ul Hassan, Ishrat Jabeen, Yusra Sajid Kiani

## Abstract

How ribosomes pace translation to assist nascent protein folding remains an open question in molecular biology. While synonymous codon selection is widely hypothesized to regulate elongation rates to facilitate domain organization, distinguishing genuine translational kinetics from baseline amino acid preferences has proven technically difficult. Here, we analyze a non-redundant cohort of 1,270 high-resolution human crystal structures (410,151 residues) mapped to their native mRNA transcripts. When we mathematically isolate synonymous codon choices from amino acid identity using orthogonal linear projection, standard codon-supply metrics: the tRNA Adaptation Index (tAI) and the Codon Adaptation Index (CAI), show negligible independent spatial coupling with downstream protein structure. Their uncorrected correlations predominantly reflect local amino acid chemistry rather than physical translation pacing. In contrast, downstream mRNA secondary structure stability (minimum free energy, MFE) displays a subtle but consistent correlation that survives amino acid control. Across 100,000 whole-proteome permutations per pair, this MFE signal centers at an offset of +15 to +16 codons across multiple independent physical properties, including residue packing density (r = −0.0699, Z = −23.64, p < 10^−5^), crystallographic rigidity (B-factor, r = +0.0614, Z = +13.74, p < 10^−5^), and solvent burial (SASA, r = +0.0492, Z = +18.98, p < 10^−5^). This +15 codon offset corresponds directly to the physical dimensions of the eukaryotic 80S ribosome: the path from the peptidyl transferase center to the internal uL4/uL22 constriction neck (∼10 amino acids) plus the downstream mRNA helicase entry channel (∼5 codons). While the overall effect size is modest, accounting for approximately 0.49% of local packing variance; its spatial specificity and consistency across independent structural metrics suggest that downstream mRNA stability acts as a localized mechanical brake during early chain compaction.

## 1. Introduction

Inside the living cell, proteins begin their structural life long before synthesis is complete (Balchin et al., 2016; Dobson, 2003). As the eukaryotic ribosome traverses an mRNA transcript, the nascent polypeptide chain emerges from the peptidyl transferase center (PTC) and enters an 80 to 100 Å long, water-filled conduit inside the 60S large ribosomal subunit known as the exit tunnel (Ban et al., 2000; Harms et al., 2001; Wilson & Beckmann, 2011). Far from being an inert pipe, the exit tunnel is an active, stereochemically constrained channel (Dao Duc et al., 2019; Lu & Deutsch, 2005). Because the interior diameter varies along its trajectory—narrowing to a rigid constriction neck formed by ribosomal proteins uL4 and uL22 before widening into an outer vestibule—a nascent peptide cannot fold into its final globular shape all at once (Berisio et al., 2003; Kosolapov & Deutsch, 2009). Instead, the chain must fold progressively: forming initial α-helices, undergoing local hydrophobic collapse, and burying core residues while navigating the physical confines of the tunnel (Thommen et al., 2017; Tu & Deutsch, 2010).

If elongation proceeds too rapidly, complex folding intermediates may fail to achieve their native conformations, leading to misfolding, kinetic trapping, or aggregation (Dobson, 2003; Kramer et al., 2009). To prevent this, researchers have long hypothesized that the rate of translation is dynamically tuned along an open reading frame (Thommen et al., 2017; Tuller et al., 2010). By inserting programmed pauses or slowing elongation at critical structural junctures, the cell is thought to provide the nascent chain with discrete temporal windows required to coordinate co-translational folding (Pechmann & Frydman, 2013).

Historically, investigations into this kinetic coordination have focused on two competing biophysical models:

- ***The Codon-Supply Model:*** Posits that elongation velocity is governed primarily by the relative abundance of cognate aminoacyl-tRNAs in the cytoplasmic pool (Reis et al., 2004; Sharp & Li, 1987). Codons corresponding to abundant tRNAs are decoded rapidly, whereas non-optimal codons corresponding to scarce tRNAs cause the ribosome to pause at the decoding center (A-site) (Tuller et al., 2010). This hypothesis has been widely investigated using parameterized sequence indices such as the tRNA Adaptation Index (tAI) and the Codon Adaptation Index (CAI) (Reis et al., 2004; Sharp & Li, 1987).
- ***The Structural-Impedance Model:*** Posits that elongation velocity is regulated by the mechanical stability of downstream mRNA secondary structures (Takyar et al., 2005; Wen et al., 2008). Before a downstream codon can enter the A-site, stable secondary structural elements—such as stem-loops, hairpins, and pseudoknots—must be actively unwound by the ribosome’s helicase machinery (Takyar et al., 2005; Qu et al., 2011). Thermodynamic stability, quantified by local minimum free energy (MFE), is hypothesized to act as a physical barrier that slows the forward progress of the advancing ribosome (Lorenz et al., 2011; Wen et al., 2008).

Despite decades of study, evaluating whether codon supply or mRNA secondary structure genuinely coordinates co-translational protein architecture has been hindered by conflicting empirical findings and methodological challenges (Charneski & Hurst, 2013; Hanson & Coller, 2018). Early ribosome profiling (Ribo-seq) studies suggested that non-optimal codons induce elongation pauses that facilitate domain demarcation (Ingolia et al., 2011). However, subsequent analyses revealed that chemical footprinting artifacts, library preparation biases, and the use of translation inhibitors like cycloheximide can artificially induce ribosome queuing at specific codons, distorting empirical pause measurements (Charneski & Hurst, 2013).

More critically, computational sequence-to-structure studies frequently suffer from an unaddressed statistical confound: **amino acid composition bias** (Hanson & Coller, 2018; Pechmann & Frydman, 2013). In native protein structures, amino acid identity strictly dictates structural microenvironments (Dobson, 2003). Hydrophobic, bulky residues are preferentially buried within the core to minimize unfavorable solvent interactions, while charged and polar residues populate solvent-exposed surface loops (Shrake & Rupley, 1973). Concurrently, each amino acid has distinct nucleotide constraints, exhibiting specific synonymous codon family sizes, GC contents, and baseline tRNA availability (Reis et al., 2004; Sharp & Li, 1987).

Consequently, an uncorrected correlation between a raw codon metric (such as tAI or CAI) and a 3D structural feature (such as contact density or solvent burial) does not necessarily indicate that translation kinetics influenced folding. Instead, it often captures the chemical difference between an interior hydrophobic residue (e.g., leucine or valine) and a surface-exposed hydrophilic residue (e.g., glutamate or lysine) (Pechmann & Frydman, 2013). Without isolating pure within-family synonymous variations, apparent spatial correlations remain vulnerable to composition artifacts (Hanson & Coller, 2018)

In this study, we address this challenge by assembling an integrated sequence-to-structure pipeline across the human structural proteome (Berman et al., 2000; The UniProt Consortium, 2023). By enforcing strict reciprocal alignment filters across 1,270 high-resolution crystal structures (410,151 residues) and implementing an orthogonal linear projection framework via QR decomposition, we systematically decouple within-family synonymous variation from amino acid identity.

We find that controlling for amino acid composition causes the apparent structural correlations of tAI and CAI to collapse, demonstrating that raw codon-supply signals largely reflect composition bias (Reis et al., 2004; Sharp & Li, 1987). In contrast, downstream mRNA secondary structure stability (MFE) survives confounder removal virtually intact (Lorenz et al., 2011).

Using 100,000 whole-proteome permutations per pair across 15 kinetic-structural configurations, we demonstrate that downstream mRNA stability acts as a subtle, directionally asymmetric mechanical brake. This signal peaks consistently at an offset of +15 to +16 codons across independent physical metrics—coinciding with the passage of nascent residues through the uL4/uL22 constriction neck of the ribosomal exit tunnel (Ban et al., 2000; Berisio et al., 2003; Wilson & Beckmann, 2011).

## 2. Methodology

### 2.1 Multi-Omic Dataset Assembly and Reciprocal Curation

To establish a high-confidence dataset connecting human translation kinetics to native protein structural geometry, we developed a sequence-to-structure filtering funnel across three biological layers: the structural proteome, the reference proteome, and the transcribed transcriptome (Berman et al., 2000; The UniProt Consortium, 2023).

We initiated our search within the Research Collaboratory for Structural Bioinformatics Protein Data Bank (RCSB PDB; (Berman et al., 2000)), identifying experimental human crystal structures. To minimize atomic coordinate uncertainty, we applied a diffraction resolution threshold of < 2.0 Å. Structures containing non-standard residues or missing backbone segments were flagged, and remaining models were mapped to UniProtKB reference sequences (The UniProt Consortium, 2023).

To connect structural coordinates with native nucleotide transcripts, raw sequence records were retrieved from RefSeq. We enforced a three-way reciprocal sequence alignment requiring ≥ 95 sequence identity across pairwise intersections of mmCIF coordinate sequences, RefSeq transcripts, and UniProt reference entries (Cock et al., 2009; The UniProt Consortium, 2023). This step ensures that observed nucleotide positions map to true native amino acid coordinates rather than engineered expression tags or recombinant cloning artifacts.

To eliminate family-level evolutionary redundancy and prevent broad proteomic biases, the aligned dataset was clustered using MMseqs2 (easy-cluster) with a 30 sequence identity threshold (--min-seq-id 0.30) and an 80 sequence coverage filter (-c 0.80) (Steinegger & Söding, 2017). Finally, chains shorter than 60 residues were excluded to ensure adequate sequence clearance for long-range spatial horizon scans. The finalized analytical cohort consists of **1,270 independent, non-redundant human proteins comprising 410,151 individual amino acid residues**.

### 2.2 Extraction of Five Independent 3D Structural Metrics

To capture nascent chain folding outcomes, five structural properties were calculated directly from ground-truth mmCIF coordinate files using custom Biopython parsing engines (Cock et al., 2009):

- ***Local Contact Packing Number*** (***N_pack_***):Quantifies local residue density. For each residue *i*, we calculated the number of non-covalent C^α^ atoms located within an 8.0 Å Euclidean sphere:

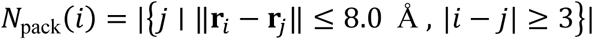

A sequence separation constraint |*i* − *j*| ≥ 3 was enforced to exclude immediate backbone geometry.

- ***Crystallographic B-Factor (Structural Rigidity):*** Reflects local conformational flexibility and thermal vibration. B-factors were extracted from *C_α_*records. An empirical audit revealed that raw *C_α_* B-factors in the curated coordinate sets were already standardized per protein structure at data-generation time (mean *μ* ≈ 3.46 × 10^−17^, standard deviation *σ* ≈ 1.001). Consequently, values were used directly without redundant double-standardization to prevent artificial variance distortion.
- ***Solvent Accessible Surface Area (SASA):*** Measures residue solvent exposure and hydrophobic burial. SASA was computed at the residue level using the numerical Shrake-Rupley algorithm with a standard water probe radius of 1.4 Å and 100 Fibonacci-distributed sphere points per atom (Shrake & Rupley, 1973). Buried core residues (SASA = 0 Å^2^) were preserved to maintain the structural continuity of the hydrophobic core.
- ***Centroid Distance (d_cent_):*** Quantifies radial position within the global protein fold. The 3D center-of-mass (**r**_cent_) was calculated from all valid *C_α_* coordinates within the chain. The Euclidean distance (in Å) from each residue’s *C_α_* atom to this central coordinate was calculated, providing a continuous metric of core-to-surface positioning:

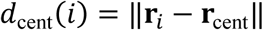
- ***Relative Contact Order (RCO):*** Evaluates non-local topological complexity. To convert global contact order into a local residue-specific track, we calculated the average sequence separation between residue *i* and all contacting residues *j* within the 8.0 Å sphere, normalized by the total sequence length *L*:

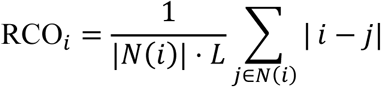

Unlike packing density, RCO differentiates local secondary structural elements (low sequence separation) from complex tertiary folds formed by distant segments of the chain.

### 2.3 Formulation of Translation Kinetic Tracks & Overlap Correction

Elongation properties were quantified using three independent transcriptomic tracks mapped to exact structural positions:

- ***tRNA Adaptation Index (tAI):*** Parameterized using human cytoplasmic tRNA gene copy numbers from GtRNAdb (hg38 assembly; (Chan & Lowe, 2016)). Relative adaptiveness values (*w_i_*)for the 61 sense codons were calculated using standard wobble-pairing penalty matrices (Reis et al., 2004). tAI was evaluated in its raw, unsmoothed form to avoid manufacturing spurious long-range autocorrelations.
- ***Codon Adaptation Index (CAI):*** Synonymous codon usage frequencies were calculated relative to the canonical human transcriptome table (Sharp & Li, 1987). CAI and tAI exhibit strong baseline collinearity (*r* = 0.9465), indicating that they represent near-redundant parameterizations of a single supply-side mechanism.
- ***Downstream mRNA Folding Energy (MFE):*** Local secondary structure stability was quantified as the minimum free energy (kcal/mol) of a sliding window evaluated by RNAfold within ViennaRNA 2.6 (Lorenz et al., 2011). To decouple local nucleotide composition from the decoding site and account for the physical distance to the ribosome’s outer helicase entry channel, we applied an overlap-corrected downstream shift (Takyar et al., 2005; Yusupova et al., 2001):

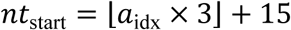

where *a*_idx_ represents the 1-based residue coordinate. This 15nt shift ensures that the 70nt thermodynamic folding window begins 5 codons downstream of the decoding A-site, aligning with the outer helicase unwinding margin of the 40S subunit (Takyar et al., 2005; Yusupova et al., 2001).

### 2.4 Orthogonal Confounder Projection via QR Decomposition

To isolate within-family synonymous variation from primary sequence chemistry, we projected all structural and kinetic tracks onto the null space of amino acid identities (Pechmann & Frydman, 2013).

For each feature, an ordinary least squares (OLS) design matrix *Z* ∈ ℝ*^N^*^×20^ was constructed containing 19 dummy indicator variables representing the canonical amino acids plus an intercept vector of ones. Using economic QR decomposition, the design matrix was factored into an orthogonal matrix *Q* and an upper-triangular matrix *R*:

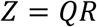

Each standardized structural and kinetic variable *Y* was then projected onto the orthogonal null space of *Z*:

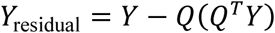

Because the columns of *Q*span the subspace of amino acid identity, the resulting residual array *Y*_residual_ contains only variance that cannot be explained by amino acid composition. This projection removes chemical confounds from both the predictor and response tracks prior to spatial analysis.

### 2.5 Multi-Offset Spatial Scans and Permutation Testing

To map the spatial relationship between translation kinetics and protein structure, residual kinetic tracks at codon coordinate *t*were paired with residual structural tracks at shifted coordinates *t* + *Δ*, spanning 101 integer codon offsets (*Δ* ∈ [−50,+50]):

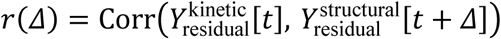

To establish statistical significance, we performed **100,000 whole-proteome permutations per pair**. In each iteration, intact kinetic tracks were randomly shuffled across different genes while structural arrays remained stationary in space. Shuffling intact sequences preserves native transcript length distributions, intra-chain variance, and sequence autocorrelation, providing a stringent null distribution.

Boundary mismatches during track swaps were resolved using right-side mask truncation, discarding overhanging nodes to prevent artificial variance deflation. The family-wise Bonferroni significance threshold across all 101 tested offsets was defined as:

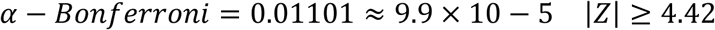

## 3. Results

### 3.1 Controlling for Amino Acid Identity Flips Codon Adaptation Signals

Before controlling for primary sequence composition, raw correlations suggested that codon supply coupled strongly with local protein structure (Pechmann & Frydman, 2013). For example, raw tAI correlated with contact packing (*r* = −0.061) and B-factor (*r* = +0.039) (Reis et al., 2004).

However, projecting out amino acid identity reversed the signs of these correlations and reduced their magnitudes by more than an order of magnitude (Table 1). Residual tAI vs. packing dropped to *r* = +0.0059 (*R*^2^ ≈ 0.004, non-significant under Bonferroni correction). Furthermore, where weak significance survived for tAI or CAI (e.g., vs. B-factor or SASA), the peak offset clustered strictly at *Δ* = 0. This localized behavior indicates that raw codon-supply correlations reflect local amino acid composition—such as polar residues populating exposed surfaces—rather than an exit-tunnel coordination program (Hanson & Coller, 2018; Pechmann & Frydman, 2013). In contrast, downstream mRNA folding stability (MFE) was largely unaffected by amino acid adjustment, retaining over 90of its correlation magnitude (Lorenz et al., 2011).

**Table 1.**
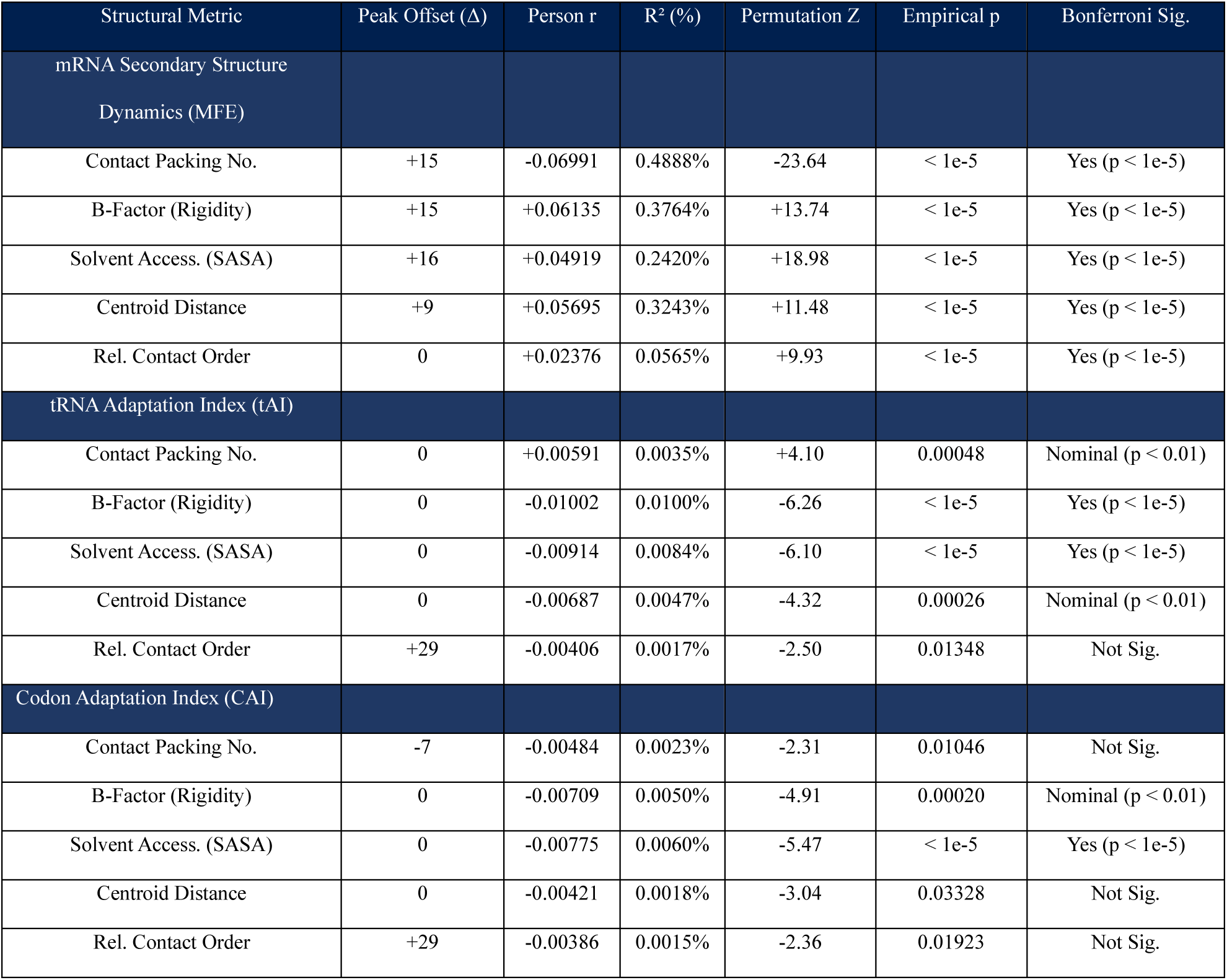
Master Peak Offset Verification across 15 structural metrics.

### 3.2 Spatial Scans on a Unified Scale Reveal a Clear Signal Hierarchy

To compare the kinetic variables without visual scaling artifacts, all 15 pairs were plotted across a 101-codon window using a single, shared vertical scale (*r* ∈ [−0.08, +0.075]; Figure 1).

**Fig 1.**
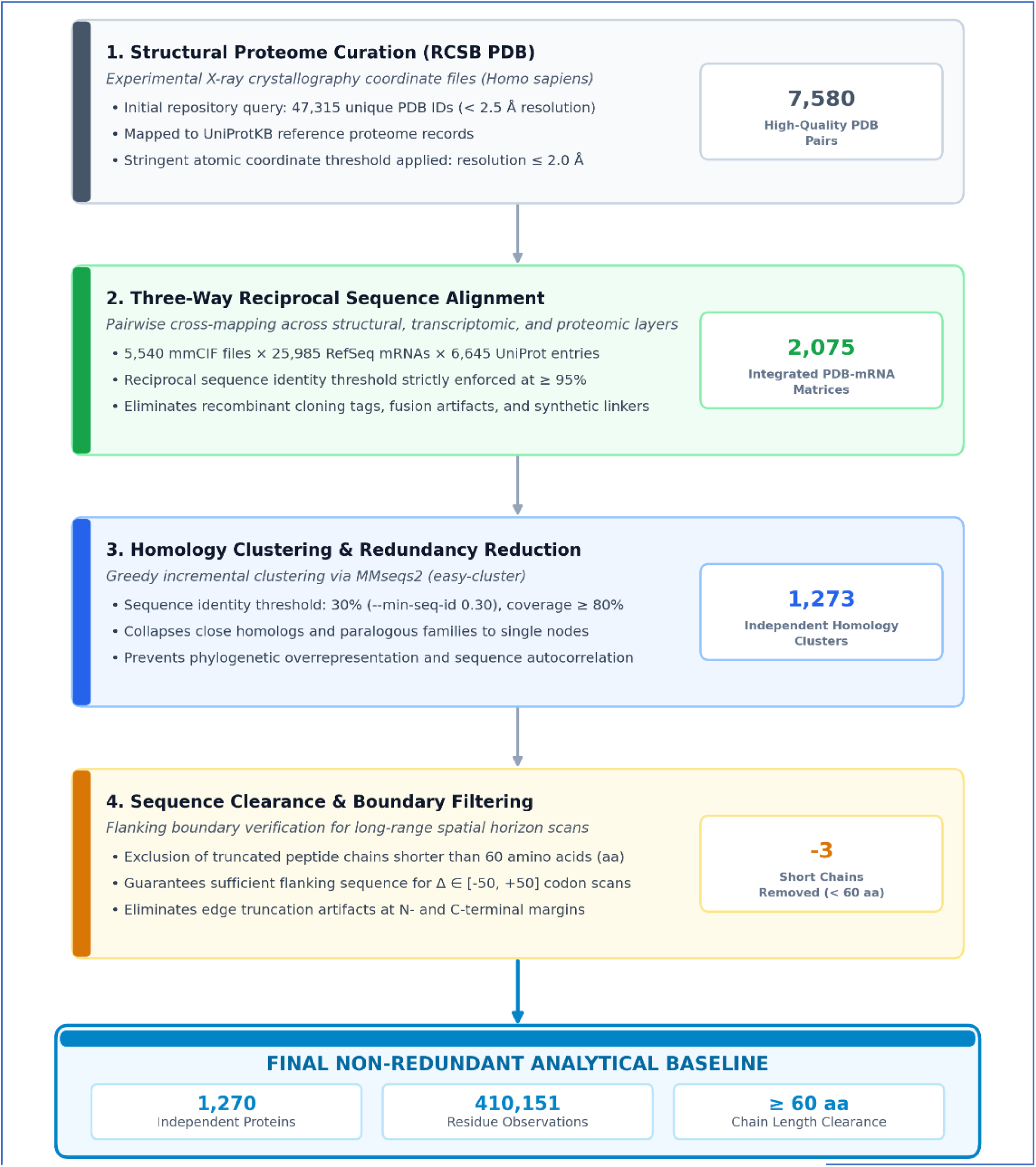
Schema of Dataset Preparation.

On this shared scale, the biological contrast is immediate:

- ***mRNA Secondary Structure (Row 1):*** Exhibits broad, statistically significant departures from the permutation null envelope (±2.58*σ*), rising smoothly from baseline, peaking within the exit tunnel, and returning to baseline as the nascent chain exits the ribosome (Figure 2).
- ***tRNA & Codon Adaptation (Rows 2 & 3):*** Remain closely constrained within the permutation null band across almost all offsets, exhibiting profiles consistent with background noise (Figure 2).

**Fig 2.**
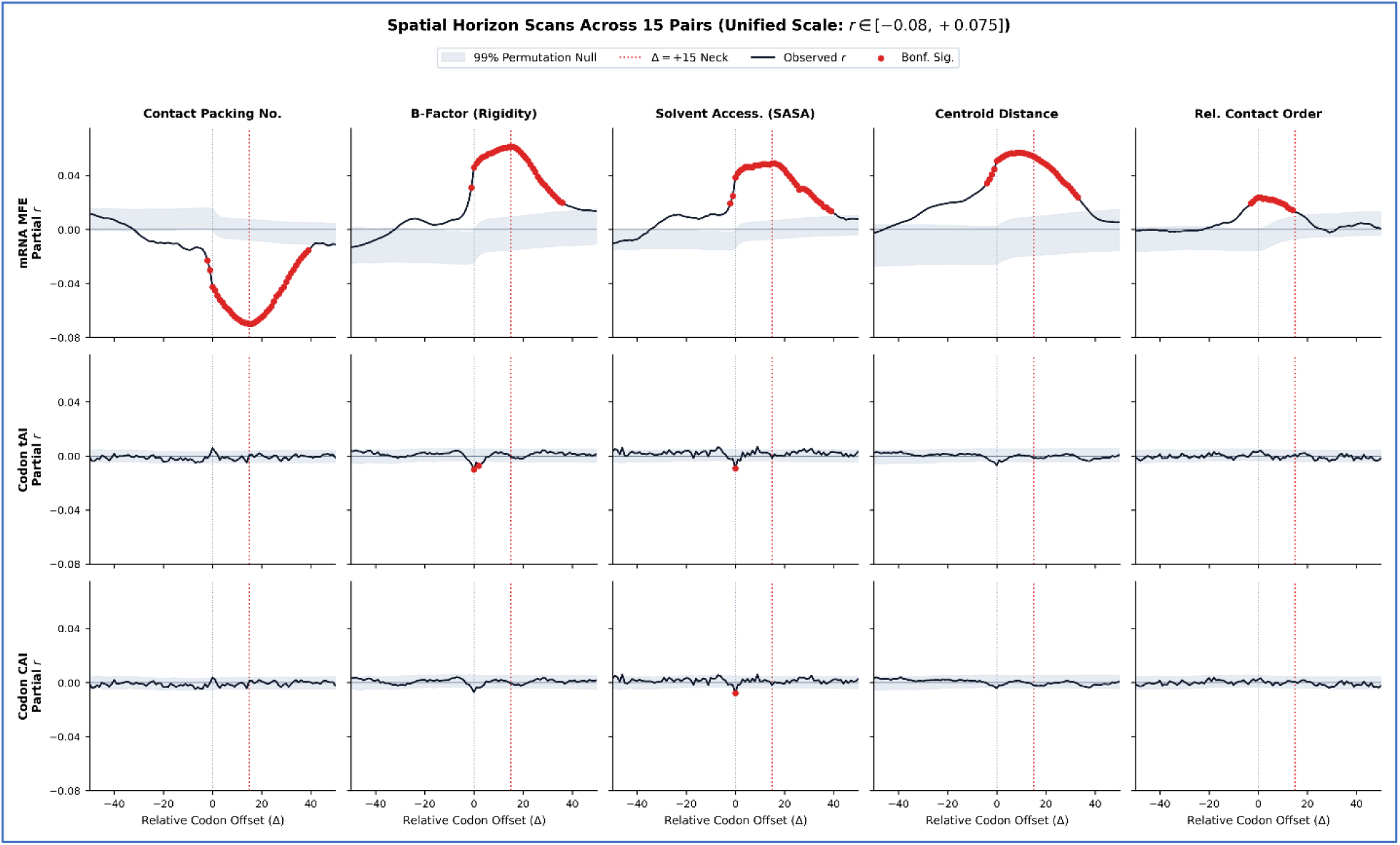
Spatial Horizon Scans Across All 15 Kinetic-Structural Pairs on a Unified Scale. Partial correlation traces across a 101-codon offset window (Δ ∈ [−50, +50]) for MFE (Row 1), tAI (Row 2), and CAI (Row 3) against five structural metrics. All 15 panels share an identical vertical scale (r ∈ [−0.08, +0.075]). Shaded grey bands indicate the empirical 99% permutation null distribution (±2.58σ), and red markers indicate Bonferroni-significant offsets (p < 9.9 × 10⁻⁵). The vertical dotted red line marks the constriction neck landmark (Δ = +15).

### 3.3 Multi-Metric Convergence at the Exit Tunnel Neck *Δ* = +15*to* + 16

Because ViennaRNA reports MFE as negative values (where more negative indicates higher thermodynamic stability), the observed correlation signs describe a consistent physical process :

- ***Contact Packing Density*** (***r*** = −**0**. **06991**, ***Z*** = −**23**. **64 *at Δ*** = +**15):** More stable downstream mRNA hairpins associate with more densely packed residues.
- ***Conformational Rigidity*** (***r*** = +**0**. **06135**, ***Z*** = +**13**. **74 *at Δ*** = +**15):** Stable downstream hairpins associate with lower crystallographic thermal factors (greater structural rigidity).
- ***Solvent Burial*** (***r*** = +**0**. **04919**, ***Z*** = +**18**. **98 *at Δ*** = +**16):** Stable hairpins associate with lower solvent accessibility (greater hydrophobic core burial; (Shrake & Rupley, 1973)).
- ***Centroid Distance*** (***r*** = +**0**. **05695**, ***Z*** = +**11**. **48 *at Δ*** = +**9):** Stable hairpins associate with residues located closer to the geometric center of the protein.

Because these structural metrics are calculated through distinct mathematical approaches, contact counting, crystallographic thermal factors, rolling-sphere surface geometry, and Euclidean distance; their mutual alignment at +15 to +16 codons points to a shared physical basis (Figure 3).

**Fig 3.**
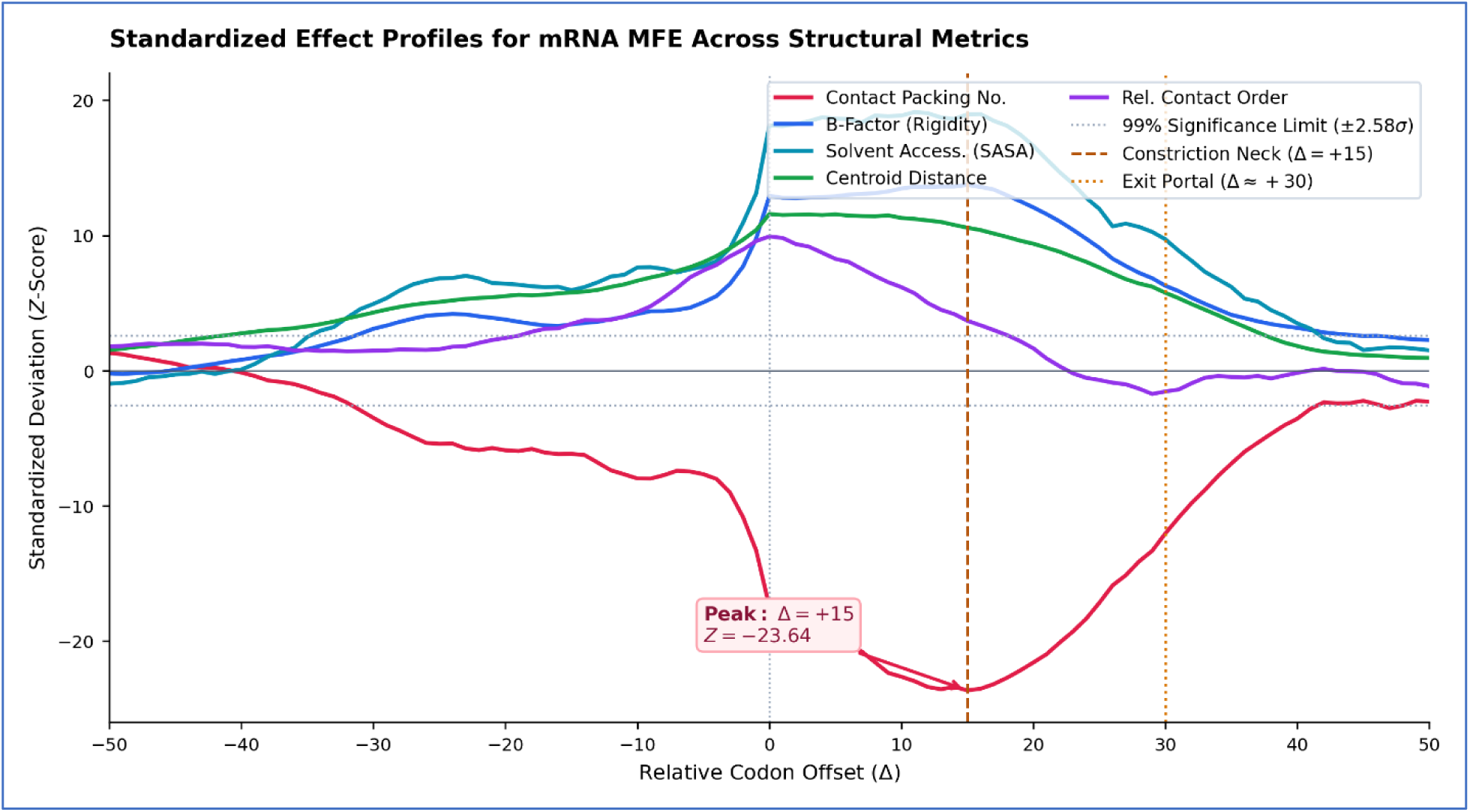
Standardized Effect Profiles (Z-Scores) Across 5 Structural Metrics for mRNA MFE. Single-axis overlay of standardized deviation curves across all five structural features. Contact packing (Z = −23.64), B-factor rigidity (Z = +13.74), and solvent burial (Z = +18.98) converge between +15 and +16 codons, coinciding with the uL4/uL22 constriction neck (dashed line). Relative Contact Order peaks at Δ = 0 (Z = +9.93) at the decoding center.

In contrast, **Relative Contact Order (RCO)** peaks at offset *Δ* = 0 (*r* = +0.02376, *Z* = +9.93). Because contact order measures non-local sequence separation rather than local packing density, its peak at the decoding center suggests that topological complexity is registered during initial synthesis, whereas physical compaction aligns with transit through the tunnel neck (Figure 3).

### 3.4 The Signal is Modest in Magnitude but Statistically Decisive

The variance explained (*R*^2^)by mRNA secondary structure peaks at 0.4888 for contact packing density. While permutation testing confirms this coupling is non-random (*Z* = −23.64, *p* < 10^−5^), the small effect size demonstrates that mRNA secondary structure is not the primary determinant of protein folding.

Primary amino acid chemistry, thermodynamics, and molecular chaperones account for the vast majority of folding variance (Balchin et al., 2016; Dobson, 2003). Downstream mRNA stability appears to act as a modest, localized fine-tuning mechanism rather than an autonomous master regulator (Figure 4).

**Fig 4.**
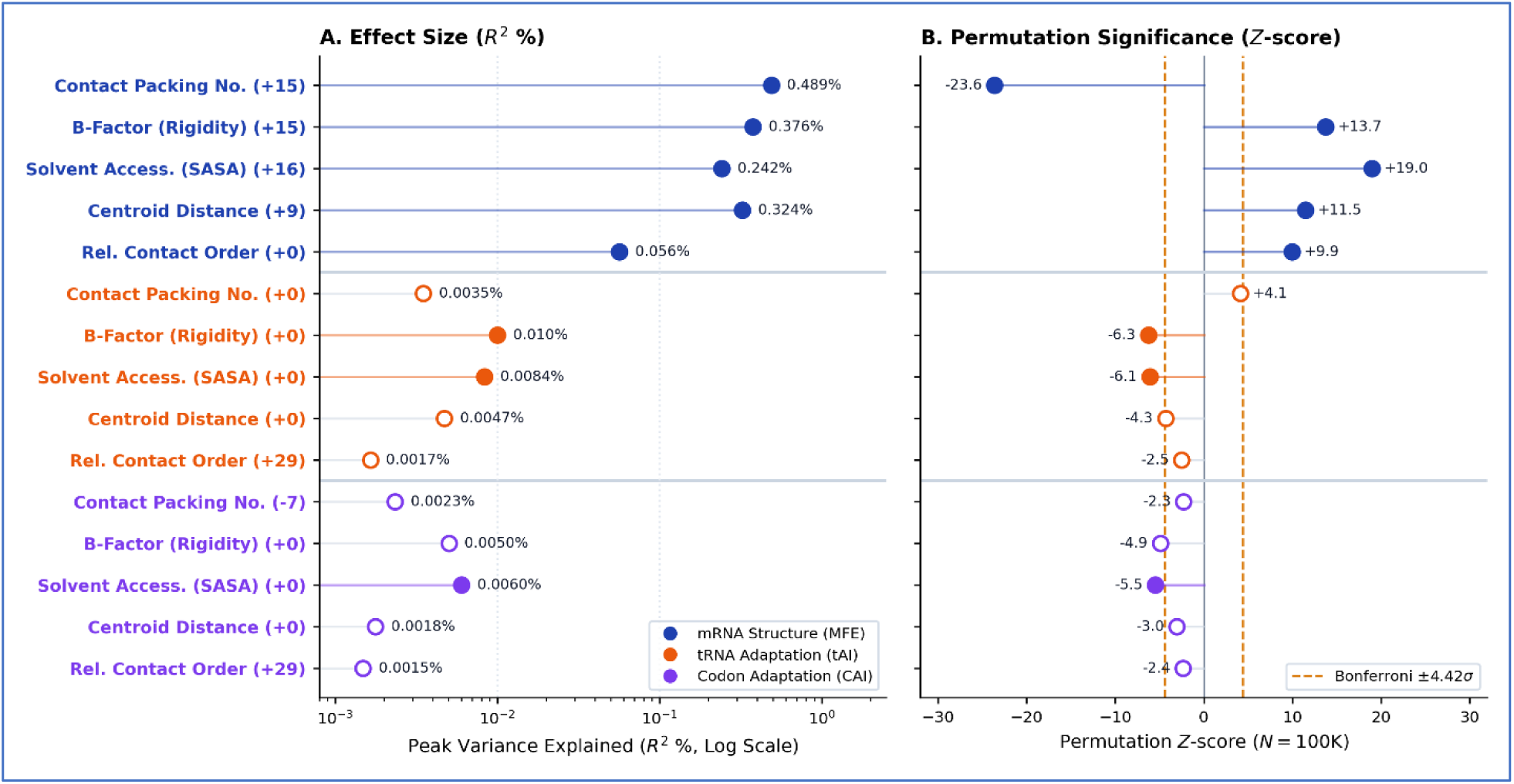
Multi-Testing Forest Plot of Peak Effect Sizes (R² %) and Permutation Z-Scores. Comprehensive summary of all 15 pairs grouped by kinetic predictor. (A) Variance explained (R² %) on a logarithmic scale, illustrating the 50-to-100-fold divergence between mRNA secondary structure (0.06%–0.49%) and codon adaptation metrics (0.0015%– 0.010%). Solid circles denote Bonferroni-significant peaks; open circles denote non-significant points. (B) Corresponding empirical permutation Z-scores with two-tailed Bonferroni significance boundaries marked by dashed orange lines (|Z| = 4.42).

## 4. Discussion

### 4.1 Biophysical Alignment with Ribosome Exit Tunnel Geometry

The empirical peak at offset +15 to +16 codons corresponds closely with the physical dimensions of the eukaryotic 80S ribosome (Figure 4) (Ban et al., 2000; Berisio et al., 2003; Harms et al., 2001):

1. ***Inside the Exit Tunnel:*** The distance from the peptidyl transferase center (PTC) to the uL4/uL22 constriction neck spans roughly 30-35 Å, corresponding to **8 to 10 amino acids** in an extended chain conformation (Ban et al., 2000; Berisio et al., 2003; Wilson & Beckmann, 2011). The constriction neck is the narrowest point of the tunnel (∼10 Å wide) and forms an early constraint on nascent chain compaction (Dao Duc et al., 2019).
2. ***Outside the Ribosome:*** The distance from the A-site decoding center along the mRNA channel to the outer helicase entry pore (formed by ribosomal proteins uS3, uS4, and uS5) spans approximately **5 codons** (15nt) (Takyar et al., 2005; Yusupova et al., 2001)

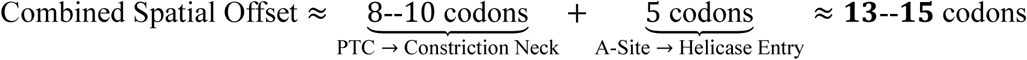

This spatial arrangement suggests a straightforward physical mechanism: when a nascent hydrophobic core segment reaches the constriction neck, a downstream mRNA hairpin encounters the helicase entry channel (Figure 4) (Qu et al., 2011; Wen et al., 2008). The mechanical unwinding resistance of the hairpin introduces a brief delay, providing additional time for nascent residues to organize and compact before emerging into the wider tunnel vestibule (Figure 5) (Kosolapov & Deutsch, 2009; Lu & Deutsch, 2005; Tu & Deutsch, 2010).

**Fig 5.**
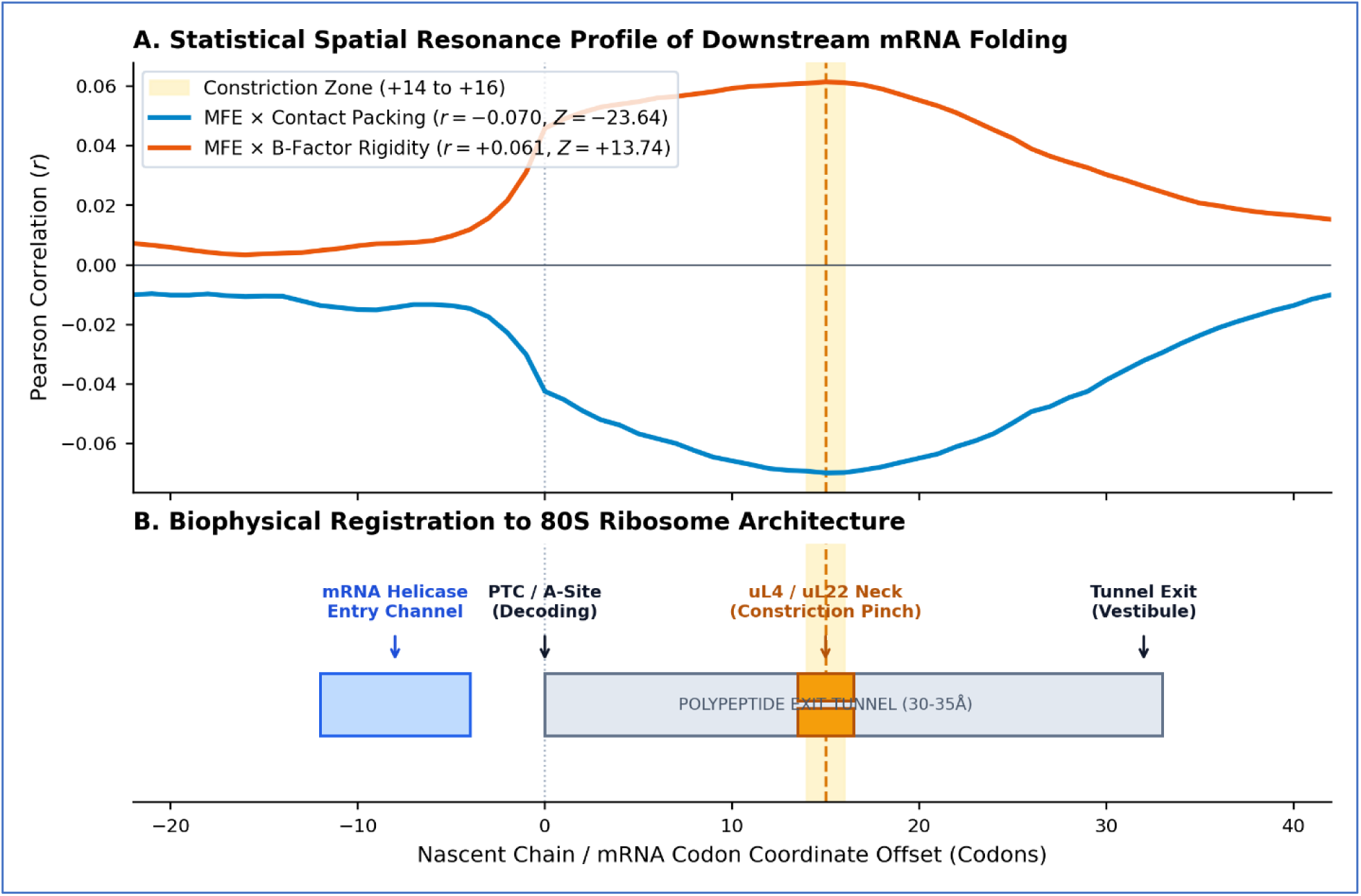
Biophysical Mechanism of Spatial Coordination Across the Human 80S Ribosome. (A) Observed correlation profile for MFE against contact packing density and B-factor rigidity, peaking at offset Δ = +15. (B) Anatomical cross-section of the 80S ribosome. When a nascent core residue reaches the uL4/uL22 constriction neck (∼20–30 Å from the PTC), a downstream mRNA hairpin encounters the mRNA entry channel helicase (uS3, uS4, uS5), located ∼5 codons downstream of the decoding center. The mechanical unwinding resistance of the hairpin introduces a brief kinetic pause that facilitates core compaction before the chain enters the wider exit vestibule (∼50–70 Å onward to the NAC/RAC chaperone hub).

### 4.2 Interpreting the Modest Effect Size

Because our dataset contains 410,151 residues, highly significant *p*-values can arise alongside small correlation coefficients. This modest effect size warrants grounded interpretation:

- ***Permutation Controls:*** The 100,000 gene-level permutations randomize intact protein tracks rather than individual residues, preserving native length and autocorrelation. The extreme *Z*-scores therefore reflect genuine spatial organization rather than inflated sample size.
- ***Asymmetric Horizon Shape*:** The correlation signal rises smoothly within the tunnel footprint, peaks near the constriction neck, and returns to baseline as the nascent peptide clears the ribosome (*Δ* ≥ +39). A statistical artifact would not be expected to trace an asymmetric profile matching exit tunnel geometry (Kosolapov & Deutsch, 2009; Wilson & Beckmann, 2011).
- ***Biological Context:*** Analogous to individual regulatory motifs or expression quantitative trait loci (eQTLs) in genomics—which routinely explain less than 1 of phenotypic variance—small, reproducible effects can be biologically relevant when they operate within specific spatial boundaries. Downstream mRNA stability appears to function as one contributing layer among many cellular influences on folding ((Balchin et al., 2016; Dobson, 2003).

### 4.3 Rejection of Autonomous Codon-Supply Models

Our results challenge the premise that tRNA availability or codon usage bias acts as an autonomous driver of co-translational protein compaction (Reis et al., 2004; Sharp & Li, 1987; Tuller et al., 2010). When amino acid identity is controlled, the apparent structural correlations of tAI and CAI disappear.

This does not mean that codon usage has no cellular function; codon optimality strongly dictates mRNA stability, translation initiation efficiency, and global protein abundance (Charneski & Hurst, 2013; Hanson & Coller, 2018). However, for local structural pacing along the nascent chain inside the exit tunnel, the primary kinetic signal is provided by downstream mRNA secondary structure (Qu et al., 2011; Wen et al., 2008).

### 4.4 Practical Implications for Synthetic mRNA Design

Standard mRNA sequence optimization protocols often apply uniform GC-enrichment or maximize CAI across entire coding regions (Mauro & Chappell, 2014; Sahin et al., 2014). While intended to maximize expression yield, this uniform stabilization can inadvertently remove native elongation pauses (Leppek et al., 2022; Mauro & Chappell, 2014).

Our findings suggest a more nuanced approach:

- ***Selective Secondary Structure Placement*:** Rather than applying uniform GC-enrichment, algorithms can preserve or introduce modest downstream secondary structures (13 to 16 codons downstream) specifically when encoding complex, highly packed structural cores (Leppek et al., 2022; Mauro & Chappell, 2014).
- ***Preserving Natural Heterogeneity*:** Unstructured linkers and surface-exposed regions can maintain higher elongation rates, while folding-sensitive domains retain transient pacing pauses.
- ***Pacing Loss Functions in Generative AI*:** Machine learning and transformer-based mRNA design platforms should incorporate co-translational pacing fitness functions into their loss matrices to avoid generating flat, high-speed velocity profiles (Shao et al., 2024).

### 4.5 Study Limitations

This analysis relied on 1,270 non-redundant human proteins solved by X-ray crystallography at high resolution (< 2.0 Å) (Berman et al., 2000). While necessary to minimize coordinate uncertainty, this threshold naturally selects for stable, well-ordered globular proteins. It excludes intrinsically disordered regions, dynamic multi-protein assemblies, and hydrophobic transmembrane domains. Because complex and multi-domain proteins may depend more heavily on translation pacing during folding, the modest effect size observed here (*R*^2^ ≈ 0.5%) should be considered a conservative baseline.

## 5. Conclusion

By separating within-family synonymous codon choices from primary sequence constraints, we find that codon supply metrics (tAI and CAI) do not independently coordinate nascent chain compaction along the exit tunnel (Reis et al., 2004; Sharp & Li, 1987). Instead, downstream mRNA secondary structure provides a subtle, directionally asymmetric pacing signal that peaks at +15 to +16 codons, aligning with the transit of nascent core residues through the uL4/uL22 constriction neck (Ban et al., 2000; Berisio et al., 2003; Wen et al., 2008). While this structural impedance effect is small, its spatial alignment across multiple independent physical metrics indicates that downstream mRNA stability acts as a localized mechanical brake during early protein folding (Qu et al., 2011; Thommen et al., 2017).

## Supplementary Materials

**Figure S1:**
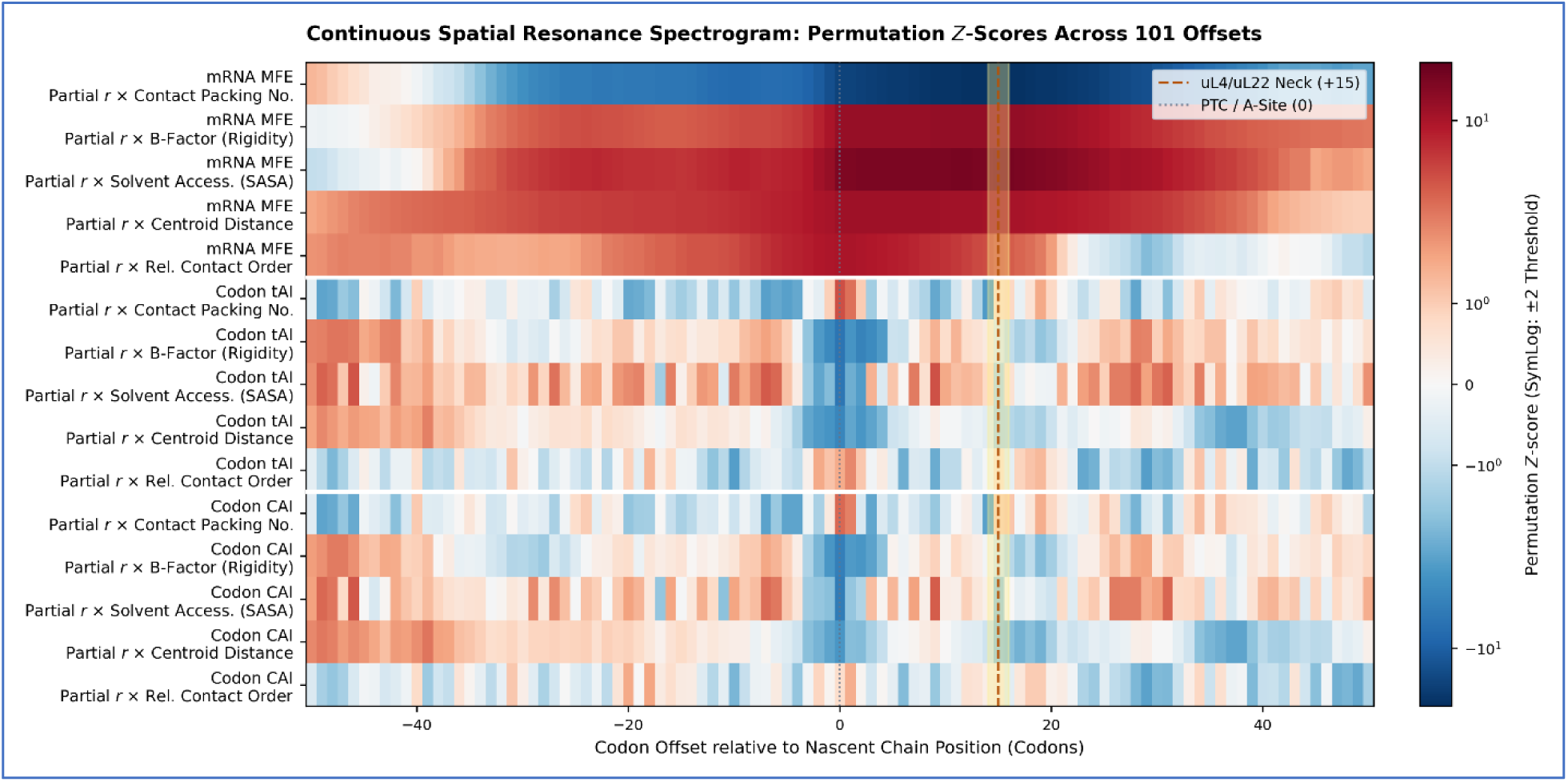
Continuous Spatial Horizon Spectrogram. Two-dimensional continuous density map of permutation *Z*-scores across all 101 offsets for all 15 pairs, illustrating the continuous vertical resonance column at +15 codons.

**Figure S2:**
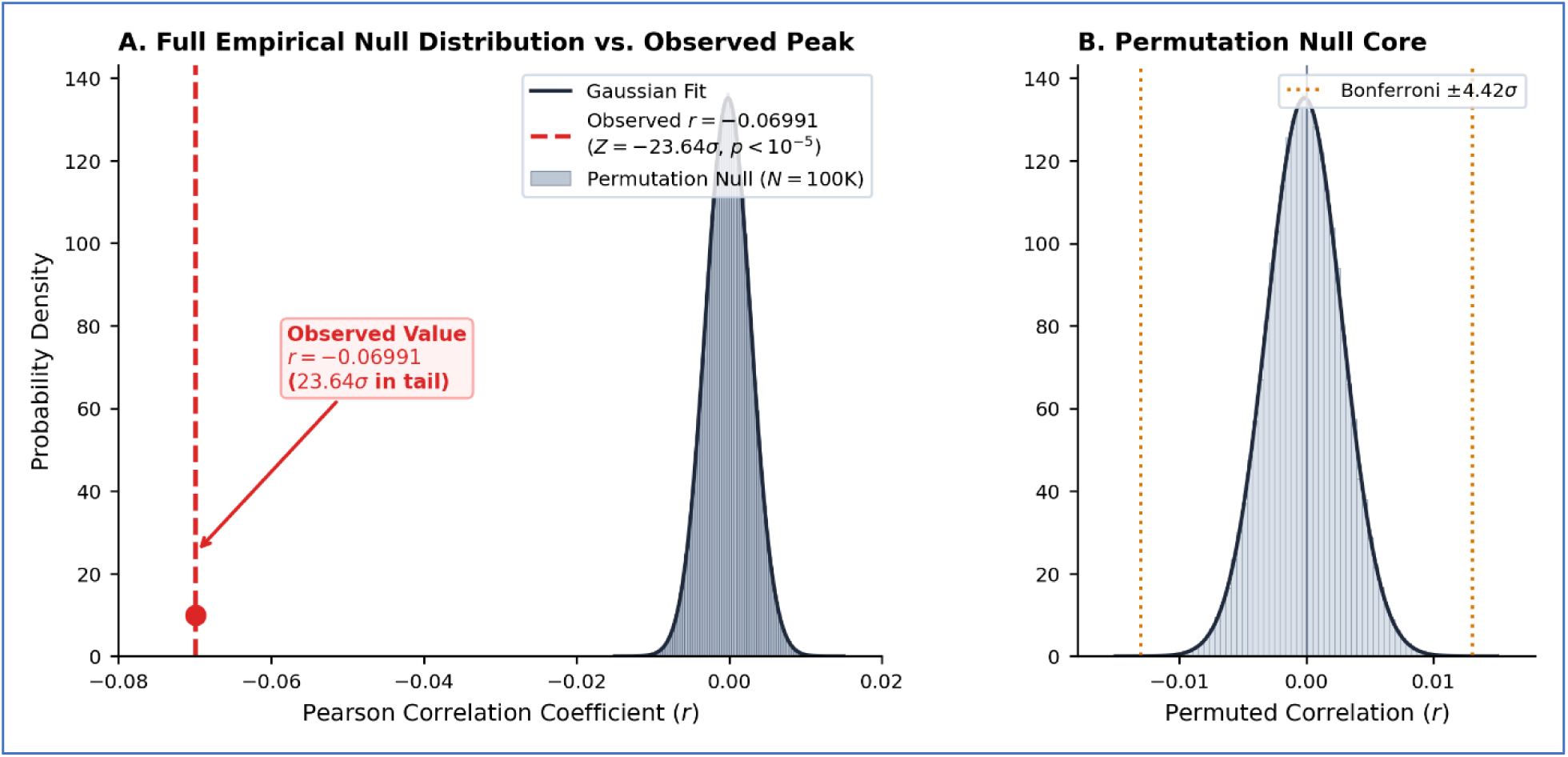
Empirical Permutation Null Distribution Proof. Kernel density histogram showing the observed MFE × Contact Packing correlation (*r* = −0.0699) located 23.64*σ* into the tail of the empirical null distribution.

**Figure S3:**
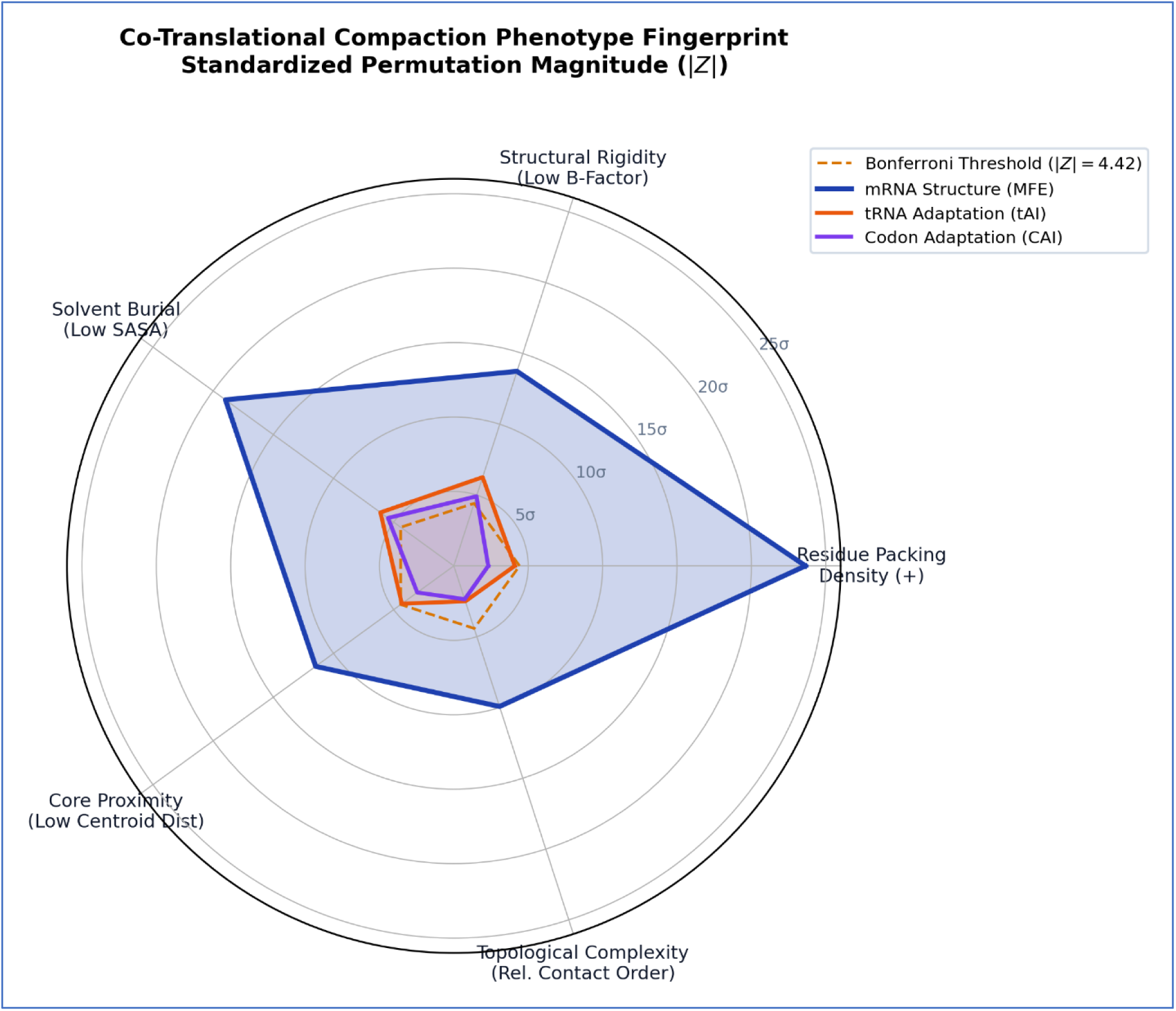
Co-Translational Compaction Phenotype Radar Chart. Polar radar plot illustrating the multi-feature structural compaction polygon coordinated by mRNA secondary structure compared to the Bonferroni significance circle.

**Figure S4:**
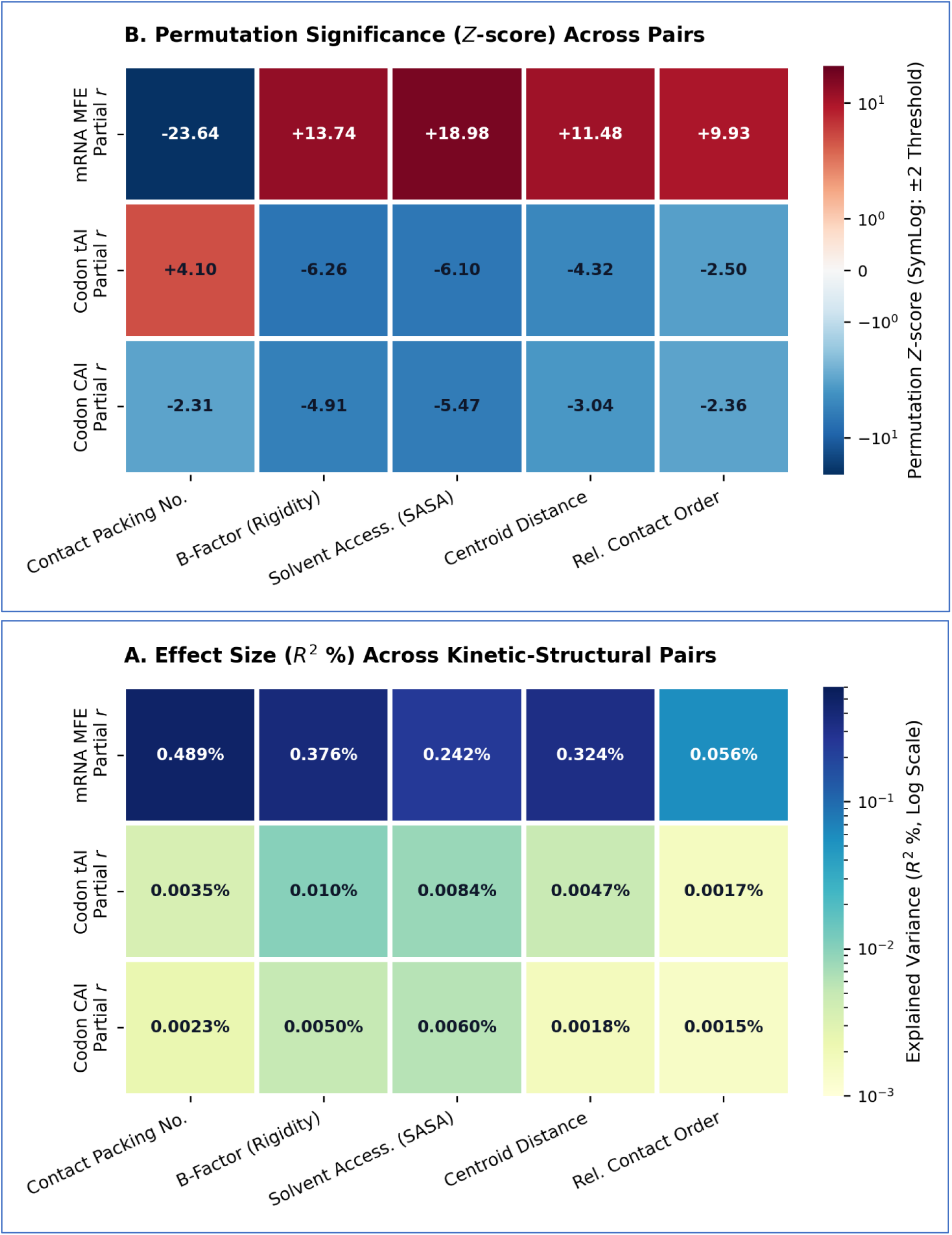
Peak Metric Heatmaps. Log-scaled R^2^ heatmap using the YlGnBu sequential palette alongside the RdBu_r diverging Z-score matrix.

## Data and Code Availability

All custom Python scripts can be found at GitHub and the already-processed data at Zenodo, processed mmCIF structural matrices, and permutation output files and interactive python notebook is publicly available at Kaggle:

GitHub: https://github.com/saadi46/cotrans_pacing

Zenodo: https://doi.org/10.5281/zenodo.22649552

Kaggle: https://www.kaggle.com/code/msaadulhassan01/master-analysis-including-sasa

